# Genomic signature of ongoing alkaline adaptation in a Schizothoracine fish (Cyprinidae) inhabiting soda lake on the Tibetan Plateau

**DOI:** 10.1101/813501

**Authors:** Chao Tong, Miao Li, Yongtao Tang, Kai Zhao

## Abstract

Comparative genomics has elucidate the molecular footprints of adaptations to extreme environments at high altitude including hypoxia, but insight into the genomic basis of saline and alkaline adaptation in highland fish has rarely been provided. The increasing of water salinization is a growing threat to Tibetan endemic fish species. Here we performed one of the first comparative genomics studies and began to characterize genomic signature of alkaline adaptation in a Schizothoracine fish inhabiting soda lake on the Tibetan Plateau. We found that expansions of lineage-specific genes associated with ion transport and transmembrane functions, genome-wide elevated rate of molecular evolution in Schizothoracine fishes relative to other lowland teleost fish species. In addition, we found specific changes in the rate of molecular evolution between *G. p. kelukehuensis* and other teleost fishes for ion transport-related genes. Furthermore, we identified a set of genes associated with ion transport and energy metabolism underwent positive selection. Using tissue-transcriptomics, we found that most REGs and PSGs in *G. p. kelukehuensis* were broadly expressed across three tissues and significantly enriched for ion transport functions. Finally, we identified a set of ion transport-related genes with evidences for both selection and co-expressed which contributed to alkaline tolerance in *G. p. kelukehuensis*. Altogether, our study identified putative genomic signature and potential candidate genes contributed to ongoing alkaline adaptation in Schizothoracine fish.

## Introduction

Being the largest and highest highland in the world with the average elevation approximately 4,000 m above sea level, the Tibetan Plateau has undergone continuous uplift during the India-Asia collision since about 45 million years ago (Li & Fang 1999; Favre et al. 2015). The uplift of the Tibetan Plateau triggered dramatic climate and environmental changes, including decreased oxygen pressure (hypoxia), long-term low temperature (chronic cold), and increased ultraviolet radiation (high UV), that imposes extremely inhospitable environmental challenges to all the wildlife dwelt at high altitude (Scheinfeldt & Tishkoff 2010; An et al. 2001). Large numbers of endemic Tibetan species had developed unique morphological, physiological or genetic features to tolerate harsh living conditions (Wen 2014). Recent studies employing genome-wide approaches mainly focused on the hypoxia and metabolic adaptation of Tibetan terrestrial animals, such as wild yak (Qiu et al. 2012), ground tit (Qu et al. 2013), and Tibetan snake (Li et al. 2018). Nevertheless, the draft genomes of very few Tibetan aquatic wildlife are sequenced (Liu et al. 2019; Yang et al. 2019), the genomic basis of aquatic animals adaptation to harsh water environments on the Tibetan Plateau still remain largely unknown. Schizothoracine fishes (Teleostei: Cyprinidae), the predominant fish fauna on the Tibetan Plateau, evolved specific phenotypic characteristics to adapt to the harsh aquatic environments, such as hypoxia, chronic cold, high UV, high salinity and alkalinity. Unveiling the genetic foundation of Schizothoracine fishes will shed novel lights on the highland adaptation of Tibetan aquatic organisms.

With the recent advances in sequencing technologies for non-model organisms without reference genomes, transcriptome sequencing is a rapid and effective approach to obtain massive protein-coding genes and molecular markers for comparative genomics (Grabherr et al. 2011). Recent comparative genomics studies based on transcriptomic data of several Schizothoracine species have identified a number of genes that underwent positive selection during the long-term adaptive processes to harsh environments on the Tibetan Plateau, such as hypoxia (Yang et al. 2014) and low temperature (Tong, Tian, et al. 2017; Tong, Fei, et al. 2017). Among these adaptive processes, evidences suggested that genes showing signals of positive selection and expansion were significantly enriched in hypoxia-inducible factor (HIF) and energy metabolic pathways. For instance, hypoxia-related genes, such as Hypoxia-inducible factor (HIF) (Guan et al. 2014) and Erythropoietin (EPO) have experienced strongly positive selection and are significantly associated with upregulation of expressions respond to hypoxia stress in Schizothoracine fish (Xu et al. 2016). In addition, genes associated with energy metabolism function, such as ATP synthase F1 subunit gamma (ATP5F1C) and ATP synthase subunit beta (ATP5b), had been detected the signature of positive selection in Tibetan naked carp that dwelt in Lake Qinghai with long-term of low temperature (Tong, Fei, et al. 2017). The main focus of the genetic mechanism of highland adaptation in Tibetan fish are still on hypoxia and chronic cold response (Yang et al. 2014; Wang et al. 2015; Kang et al. 2017; Guan et al. 2014; Xu et al. 2016). Notably, an increasing number of lakes on the Tibetan Plateau are existing or towards saline and alkaline due to the global climate changes and human activities (Zheng 1997; Yang et al. 2010). The Tibetan freshwater endemic fishes are long suffering these harsh conditions challenges. However, the genomic signature of high salinity and alkalinity adaptation in Schizothoracine fish have yet to be comprehensively determined. Therefore, it may provide novel insights for understanding the mechanism of highland adaptation of Tibetan fish.

A Schizothoracine fish, *Gymnocypris przewalskii kelukehuensis* (Cyprinidae) only inhabited in soda lake located at the Tsaidam Basin on the northestern Tibetan Plateau (Tong, Tang, et al. 2017) (FIG. 1A and 1B). Lake Keluke is a brackish-water lake with the salinity of 0.79‰ and pH value up to 8.5, while Lake Tuosu is a typical salt lake with unusually high sodium, potassium and magnesium concentration, its salinity even reaches 26.5‰ (FIG. 1C) (Zheng 1997). Both lakes are connected and alkaline lake Tuosu used to be freshwater. Notably, Lake Tuoso dramatically evolved to be typically salt water with extremely high salinity, while Lake Keluke are gradually evolving towards high salinity and alkalinity (Wu 1992; Zheng 1997). Past evidence showed that *G. p. kelukehuensis* used to inhabit in both lakes, while it is now rare in Lake Tuosu. This indicated that *G. p. kelukehuensis* mainly dwelt in Lake Keluke and was suffering the challenge of living aquatic environment towards high salinity and alkalinity. Because of the unique evolutionary history on Tibetan Plateau at high altitude, *G. p. przewalskii* provides an exceptional model to investigate the genetic mechanisms underlying ongoing alkaline adaptation on the Tibetan Plateau. In addition, human activities and exotic species invasions could drive the declines of biodiversity in remaining intact ecosystem fragments and accelerating species extinctions (Ceballos et al. 2015). Recent evidences revealed that overfishing and exotic species invasions (*Cyprinus carpio, Hypophthalmichthys molitrix* and *Ctenopharyngodon idella*) had dramatically deceased the natural population size of *G. p. kelukehuensis* (Wu 1992; Zhang et al. 2013; Tong, Tang, et al. 2017). Thus, characterization of its genomic resource will contribute to its conservation.

**FIG. 1.**
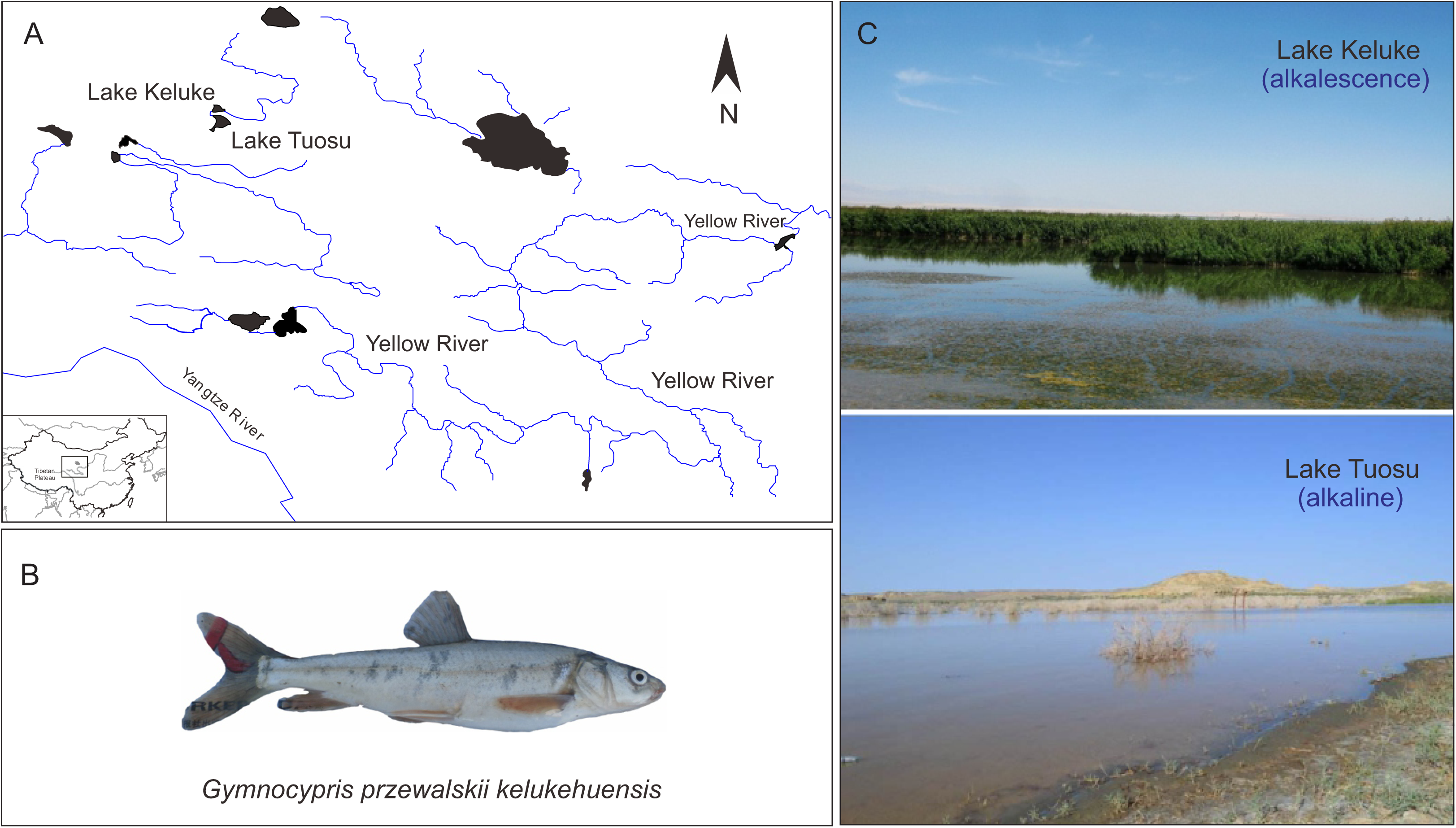
The sampling site and alkaline lake on the northeastern Tibetan Plateau. (A) The sampling map of *Gymnocypris przewalskii kelukehuensis* in Lake Keluke at the Tsaidam Basin on the northwestern Tibetan Plateau. (B) Representative specimens of *G. p. kelukehuensis*. (C) Representative alkaline lake at the Tsaidam Basin, including Lake Keluke (top) and Lake Tuosu (bottom). Photo credit: Chao Tong.

In this study, we generated the first transcriptome of a Schizothoracine fish, *G. p. kelukehuensis*. We focused in particular on identifying putative genomics signatures of ongoing alkaline adaptation of fishes inhabiting soda lake on the Tibetan Plateau. Specifically, we performed the comparative genomics study together with recently available Schizothoracine transcriptomes and other teleost fish genomes. We aimed to identify genome content and genome-wide pattern of molecular evolution that differ between fish species. In addition, we sought a set of genes underwent rapidly evolving and positive selection contributed to its adaptation. Moreover, using new tissue-specific expression data, we characterized the expression pattern of genes under natural selection, and predicted the coexpression network contributed to alkaline tolerance.

## Materials and Methods

### Sample collection and transcriptomics

A total of four adult *G. p. kelukehuensis* individuals (two males and two females) with similar body size (113 ± 0.9 g) were collected from Lake Keluke (37°19’17.2”N, 96°53’14.0”E) in October, 2016 using gill nets. All fish samples were dissected after anesthesia with MS-222 (Solarbio, Beijing, China). Three tissues (gills, head kidney and intestine) were collected from each individual and immediately stored in liquid nitrogen at −80 °C. All the animal experiments were approved by the Animal Care and Use Committees of the Northwest Institute of Plateau Biology, Chinese Academy of Sciences.

Sample of each tissue was marked separately, resulting in four biological replicates for each of the four tissues. Total RNA of each sample was extracted using TRIzol reagent (Invitrogen, CA, USA) in accordance with manufacturer’s instructions, and detected for quality and quantity of RNAs with Nanodrop 1000 (NanoDrop Technologies, DE, USA) and Agilent Bioanalyzer 2100 (Agilent Technologies, CA, USA). Only RNA with high quality (RNA Integrity Number > 7) were used for cDNA synthesis and amplification. Libraries were prepared with Nextera XT DNA Sample Preparation Kit (Illumina, CA, USA) using ∼350-bp inserted fragments for transcriptome sequencing as previously described (Tong, Tian, et al. 2017; Tong, Fei, et al. 2017; Tong et al. 2019). Libraries were individually barcoded and run on a single lane of an Illumina NovaSeq (Novogene, CA, USA) yielding 150-bp paired-end (PE150) reads.

Illumina sequencing reads were checked for quality using package, FastQC. Sequencing adapters and reads with a quality score < 20 were trimmed with Trimmomatic (Bolger et al. 2014), resulting in clean reads. We built a *de novo* transcriptome assembly based on clean reads using Trinity v2.6.5 (Grabherr et al. 2011) with default parameters. Next, we removed the redundant transcripts using CD-HIT (Fu et al. 2012) with the threshold of 0.90 and extracted the longest transcript as unigenes. We predicted the open reading frame (ORFs) of each transcript using TransDecoder (https://github.com/TransDecoder/TransDecoder) and MARKER (Cantarel et al. 2008). We used the BUSCO v3.0 (Simão et al. 2015) package based on Actinopterygii_odb9 gene set from OrthoDBv9 to assess the completeness of the assembly.

We also searched all published Schizothoracine fish transcriptomes (Tong, Fei, et al. 2017; Zhang et al. 2017; Chi et al. 2017) and downloaded the sequencing reads from NCBI SRA database (https://www.ncbi.nlm.nih.gov/sra) (supplementary table S1). We assembled all the download RNA-seq data following above pipeline.

### Orthologs identification

We searched and downloaded seven well-annotated teleost fish genomes from Ensembl database (release 96) (http://useast.ensembl.org/index.html) to build a local protein database, including zebrafish (*Danio rerio*), cod (*Gadus morhua*), cave fish (*Astyanax mexicanus*), fugu (*Takifugu rubripes*), tilapia (*Oreochromis niloticus*), medaka (*Oryzias latipes*) and spotted gar (*Lepisosteus oculatus*) (FIG. 2A). We translated *G. p. kelukehuensis* nucleotide sequences of protein coding genes into amino acid sequences using a custom perl script, and pooled this dataset into existing protein database. Next, we conducted the self-to-self BLASTP for all amino acid sequences with a E-value cutoff of 1e^−5^, and removed hits with identity□<□30% and coverage□<□30%. We identified the orthologous gene groups from the BLASTP results with OrthoMCL v2.0.9 (Chen et al. 2006) with default settings. We calculated, mapped and illustrated all the identified orthologous gene groups of each species by venn diagram. Finally, we identified lineage-specific orthologous gene groups in *G. p. kelukehuensis*. We carried out the Gene Ontology (GO) functional enrichment analyses for these set of genes using Blast2GO software (Conesa et al. 2005).

**FIG. 2.**
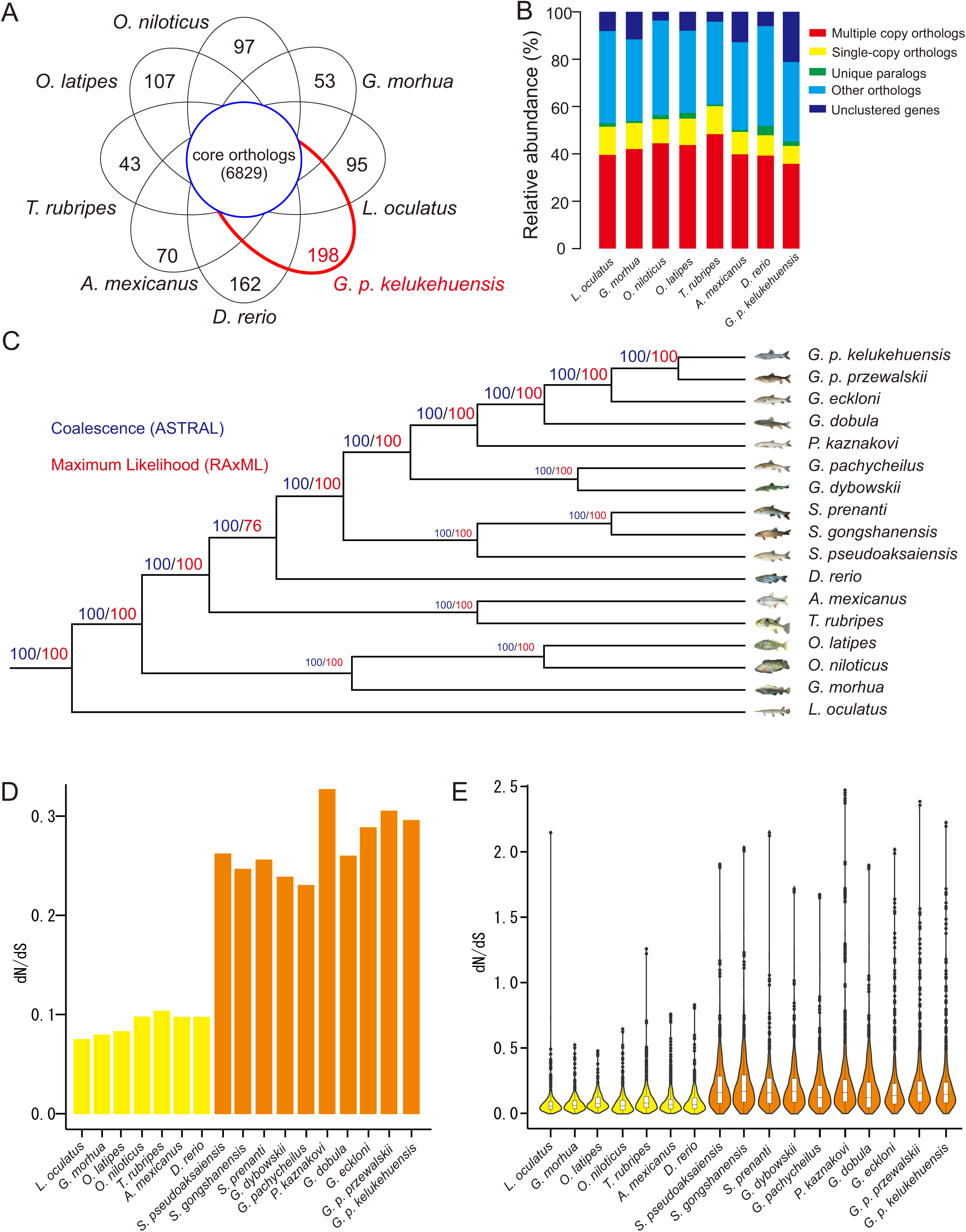
Comparative genomic features of *G. p. kelukehuensis* and other fish species. (A) Venn diagram showed shared and distinct orthologous gene groups (gene families) belonged to eight fish species. The number of core orthologous genes shared by eight species was 6,829. (B) Spine plot depicting the composition of different categories of gene families labeled by colors. (C) Genome-wide phylogeny of 17 fish species.

### Genome content comparison

Among the genes under the above identified orthologous gene groups, we identified one to one, one-to-many, and many-to-many orthologs among eight fish species. Notably, for each 1:1 ortholog pair, we only selected the longest transcript associated with the gene for each pair of species as putative single-copy orthologs. Next, the number of genes under each category of above identified orthologs in each fish species were calculated, respectively.

Using above identified shared single-copy orthologs as seed, we identified the curated orthologs in other nine Schizothoracine fish species with E-values of less than 10^−20^ by HaMStR (Ebersberger et al. 2009). We aligned and trimmed the protein sequences of all orthologous groups using PRANK (Löytynoja & Goldman 2005) with the parameter “-codon” and MATFF (https://mafft.cbrc.jp/alignment/software/), and trimmed using trimAl (Capella-Gutiérrez et al. 2009) with the parameter “-automated1”, which resulting in putative single-copy orthologs of each Schizothoracine fish for further phylogenomics.

### Genome-wide phylogeny construction

We aligned each single-copy ortholog of all fish species using MUSCLE v3.8.31 (https://www.ebi.ac.uk/Tools/msa/muscle/) with default parameters and trimmed using trimAl (Capella-Gutiérrez et al. 2009) with parameter “-automated1”. To maximize the information content of sequences and minimize the impact of missing data, we filtered the core single-copy orthologs with stricter constraints, including length (minimum 200 aa), sequence alignment (maximum missing data 50% in CDS alignments). After filtration, we prepare two types of gene datasets. At first, we concatenated all core single-copy genes of each species into one-line sequence as a supergene using a custom python script (genome-scale concatenation-based). In addition, we also conducted a genome-scale coalescent-based dataset including thousands of core single-copy genes. We detected the best model for tree construction using ModelTest2 (Posada & Crandall 1998), and then we built the Maximum Likelihood (ML) phylogenetic tree using RAxML 8 (Stamatakis 2014). Finally, we reconstructed the species tree using ASTRAL 4.4.4 (Mirarab et al. 2014).

### Nucleotide substitution rate estimation

To estimate lineage-specific evolutionary rate for each branch of the phylogeny of 17 fish species, we aligned single-copy orthologs using MUSCLE v3.8.31, derived nucleotide alignments from protein alignments using PAL2NAL v14 (Suyama et al. 2006), and estimated pairwise dN/dS of nucleotide alignments using the CodeML package in PAML 4.7a (Yang 2007). Specifically, we used the free-ratio model (“several ω ratio”) to calculate the ratio of dN to dS nucleotide changes separately for each ortholog and a concatenation of all alignments of 2,178 single-copy orthologs from the 17 fish species. Parameters, including dN, dS, dN/dS, N*dN, and S*dS values, were estimated for each branch, and genes were discarded if N*dN or S*dS < 1, or dS >2, following previous studies (Tong, Tian, et al. 2017; Tong, Fei, et al. 2017; Tong et al. 2015).

### Rapidly evolving gene identification

We used two branch models in CodeML package (Yang 2007) to identify rapidly evolving genes (REGs) in *G. p. kelukehuensis* lineage with corresponding nucleotide alignments, specifically with the null model assuming that all branches have been evolving at the same rate and the alternative model allowing the focal foreground branch (*G. p. kelukehuensis*) to evolve under a different evolutionary rate. We used a likelihood ratio test (LRT) in R software, package MASS with df□=□1 to discriminate between the alternative model and the null model for each single-copy orthologs in the genesets. We only consider the genes as evolving with a significantly faster rate in the foreground branch if the adjusted *P* value□<□0.05 and higher dN/dS in the focal foreground branch than focal background branches (other 16 fish species).

### Positively selected gene identification

We used a branch-site model in CodeML package to identify genes with positively selected amino sites (PSGs) in *G. p. kelukehuensis* lineage. Specifically, we treated *G. p. kelukehuensis* as the focal foreground branch, with the other 17 fish lineages being specified as the background branch. We conducted a LRT to compare a model that allows sites to be under positive selection (dN/dS >□1) on the foreground branch with a posterior probability more than 0.95 based on the Bayes empirical Bayes (BEB) statistic. We also calculate P values based on a Chi-square statistic adjusted by the FDR method as described above. We treated the genes with adjusted P value□<□0.05 as candidates experiencing positive selection. Finally, we annotated the Gene Ontology (GO) functional categories of both REGs and PSGs, and identified significantly enriched GO categories using R software, package topGO (Alexa & Rahnenfuhrer 2010).

### Gene expression analysis

We mapped all the clean reads to all assembled unigenes using RSEM (Li & Dewey 2011) to obtain expected counts and transcripts per million (TPM). To avoid the contamination during dissection causes the low levels of expression in the neighboring structures, we removed genes with TPM < 1 in each replicate of three tissues. To classify genes by their tissue specificity, we calculated τ, a commonly used metric of expression specificity (Yanai et al. 2005). τ ranges from 0 to 1, where 0 indicates that genes are ubiquitously expressed and 1 indicates that genes are exclusively expressed in one tissue. We focused on genes of at least moderate expression because we are primarily interested in REGs and PSGs of very high expression in one tissue that are only lowly expressed in another tissue are candidates for being false positives for expression in the second tissue.

### Construction of gene co-expression network

We performed the weighted gene co-expression network analysis (WGCNA) to predict the putative regulatory networks involved in alkaline tolerance. Specifically, we imported all the gene expression data into R package, WGCNA (Langfelder & Horvath 2008) to generate coexpression modules using the blockwiseModule function for automatic network construction with default parameters. Modules were defined as clusters of highly interconnected genes. For each module, a module eigengene (ME) was the first principal component of a cluster of genes within the module, which represented the module’s gene expression profile. Highly correlated modules (in terms of ME) were further merged using mergeCutHeight of 0.25. Briefly, the grey module included genes that cannot be classified into any other modules. In addition, we screened the genes in each modules with high and significant correlation including REGs or PSGs, and executed an additional filtration. Finally, we annotated the genes in each module by GO using R package topGO (Alexa & Rahnenfuhrer 2010).

## Results

We generated approximately 116.11 Gb raw data from the transcriptome sequencing of three tissues with four replicates for each (supplementary table S2), including gill, head kidney and intestine of a Schizothoracine fish, *G. p. kelukehuensis*. After trimming, we obtained an average 65 million PE150bp reads from each replicate. After *de novo* assembly, the transcriptome yielded 408,614 transcripts in total, with a mean length of 941 bp and N50 of 1,787 bp. After removing redundant isoforms and extraction of longest isoform among alternative transcripts, a total of 30,670 unigenes were yielded, ranged from 201 to 23463, with an N50 of 3,072 bp and an average length of 1,985 bp (supplementary table S3). In total, we obtain 28,815 protein-coding genes with full length or partial of gene coding regions (supplementary table S4). Of the 4,584 highly conserved orthologs recoded in the Actinopterygii_db9 database, we identified 4,231 (92.3%) of seed orthologs to be present in our assembly.

### Expensive lineage-specific genes of *G. p. kelukehuensis* targets in ion transport function

A total of 174,711 proteins from above eight fish species protein coding genesets were binned into 30,211 orthologous gene groups (gene family). We identified a total of 6,829 core orthologous gene groups shared by eight fish species (FIG. 2A). Besides of shared orthologous groups, we found a number of putative lineage-specific gene groups in each fish species (FIG. 2A), such as 198 gene groups identified in *G. p. kelukehuensis*. Functional enrichment analysis suggested that lineage-specific genes in *G. p. kelukehuensis* were mainly associated with ion transport and transmembrane transportation functions (supplementary table S5), such as monovalent inorganic cation transport (GO:0015672, P□=□0.000031), response to pH (GO:0009268, *P*□=□0.0072), intracellular pH reduction (GO:0051452, *P* = 0.00576), and water transport (GO:0006833, *P*□=□0.00456).

After comparing the number of orthologous gene groups among eight fish species, we found that 22,725 (78.87%) *G. p. kelukehuensis* genes clustered into 15,574 gene families (supplementary table S6). We also identified similar number of gene families in zebrafish genome, while with more orthologs than *G. p. kelukehuensis*. Notably, for the unclusted genes, *G. p. kelukehuensis* had the largest number (N = 6,092), while only the smallest number was found in *T. rubripes* genome (N = 768) (FIG. 2B).

### Phylogenomics pinpoints the phylogenetic position of *G. p. kelukehuensis* within Schizothoracinae clade

After strict 1:1 ortholog selection, we identified 2,178 putative single-copy orthologs among 6,829 shared orthologs (only one ortholog in each gene family) in each fish species. After filtration, we eventually obtained 1,159 orthologs and concatenated them into a single supergene for each fish species. We constructed two maximum likelihood phylogenetic trees of 17 fish species based on the concatenated (supergene) and coalesced single-copy orthologs using protein sequence datasets. Both combination of reconstruction methods (coalescent-based or concatenation-based) yielded the same topology (FIG. 2C). Past studies suggested that Schizothoracine fishes could be divided into three groups: primitive, specialized and highly specialized groups based on their phenotypic traits and living environment conditions. Each group represents a specific historical stage associated with the phased uplift of the Tibetan Plateau (Cao et al. 1981; Zhang et al. 2018). As genus *Gymnocypris* belonged to the highly specialized group of Schizothoracinae, it is not surprising that *G. p. kelukehuensis* and other three *Gymnocypris* species were clustered into one clade (FIG. 2C). Besides the newly reconstructed phylogeny within Schizothoracine fishes, the strongly supported (100%) phylogeny also had similar topology to recent phylogenomics studies on Schizothoracine fishes and other teleost fishes (Tong, Fei, et al. 2017; Zhang et al. 2017; Chi et al. 2017).

### Schizothoracine fishes exhibit the distinct genome-wide signature of accelerated protein evolution

We identified the signature of natural selection acting on each species based on 2,178 shared single-copy orthlogs using different models in PAML (Yang 2007). We found that Schizothoracinae clade including *G. p. kelukehuensis* lineage had an elevated genome-wide dN/dS ratio compared with other seven teleost fish species (FIG. 2D). In addition, we analyzed the dN/dS of each shared single-copy ortholog among involved fish species. Intriguingly, using both comparison strategies, we found Schizothoracinae clade exhibited an elevated patterns of nucleotide substitution rate than non-Schizothoracinae clade at a genomic scale (FIG. 2E).

### Abundant rapidly evolving genes involved in ion transport process

Accelerated evolution at molecular level may be reflected by an increased rate of non-synonymous changes within genes involved in adaptation (Zhang et al. 2014). We identified 475 putative REGs (*P* < 0.05) that had elevated rates of molecular evolution (dN/dS) in *G. p. kelukehuensis* lineage relative to other involved fish species (supplementary table S7). The most important finding was that REGs included genes functioning in ion transport and ion channel, such as solute carrier (SLC) family (SLC13A3, SLC39A9 and SLC10A2), transmembrane protein (TMEM) family (TMEM9SF3, TMEM33, TMEM97, TMEM120, TMEM175, TMEM177, TMEM208) and H^+^/Cl^-^ exchange transporter 7 (CLCN7).

### Gene associated with ion transport and energy metabolism underwent positive selection

Positive selection analysis could pinpoint genes that associated with a functional and environmental shift (Stanley 1975). We identified 66 candidate orthologs that underwent positive selection (PSGs) in *G. p. kelukehuensis* lineage. Functional enrichment analysis result showed that above identified PSGs were significantly associated with ion transport and energy metabolism functions (supplementary table S8), partially similar GO categories as REGs. We identified a set of ion transport-related PSGs, such as solute carrier organic anion transporter family member 4A1 (SLC4A1) associated with sodium-independent organic anion transport. In addition, we found a number of PSGs involved in energy metabolism, such as cAMP-dependent protein kinase catalytic subunit (PRKACA), inositol-trisphosphate 3-kinase A (ITPKA) and molybdenum cofactor biosynthesis protein 1 (MOCS1).

### Genes under selection tend to broadly-expressed in *G. p. kelukehuensis*

We focused on the expression patterns of REGs and PSGs between the three tissues associated with alkaline tolerance in *G. p. kelukehuensis*, including gills, head kidney and intestine. Most REGs (N = 395) were expressed in the head kidney, while less number of expressed REGs (N = 359) was found in the intestine (FIG. 3A, supplementary table S9). We also found the similar signature of expression patterns in PSGs, highest number (N = 53) of expressed PSG was found in head kidney and less (N = 45) in intestine (FIG. 3A, supplementary table S10). Notably, most of REGs (N = 228) and PSGs (N = 28) were broadly-expressed in all three tissues. Among the broadly-expressed REGs, three tissue types shared the similar expression signatures, such as the range of expression levels (FIG. 3B). Notably, a set of broadly expressed solute carrier and transmembrane protein genes were all identified. GO enrichment analysis showed that this set of broadly-expressed REGs was significantly enriched in ion transport function, such as sodium ion transport (GO: 0006814, *P* = 0.00021), metal ion transport (GO:0030001, *P* = 0.00018), zinc ion transport (GO:0006829, *P* = 0.00035) and chloride transmembrane transport (GO:1902476, *P* = 0.00014) (FIG. 3C). In addition, we also found a set of broadly-expressed PSG were associated with ion transport function, such as sodium-independent organic anion transport (GO:0043252, *P* = 0.00018), sodium ion transport (GO: 0006814, *P* = 0.00044) and ion transmembrane transport (GO: 0034220, *P* = 0.00036) (FIG. 3D).

**FIG. 3.**
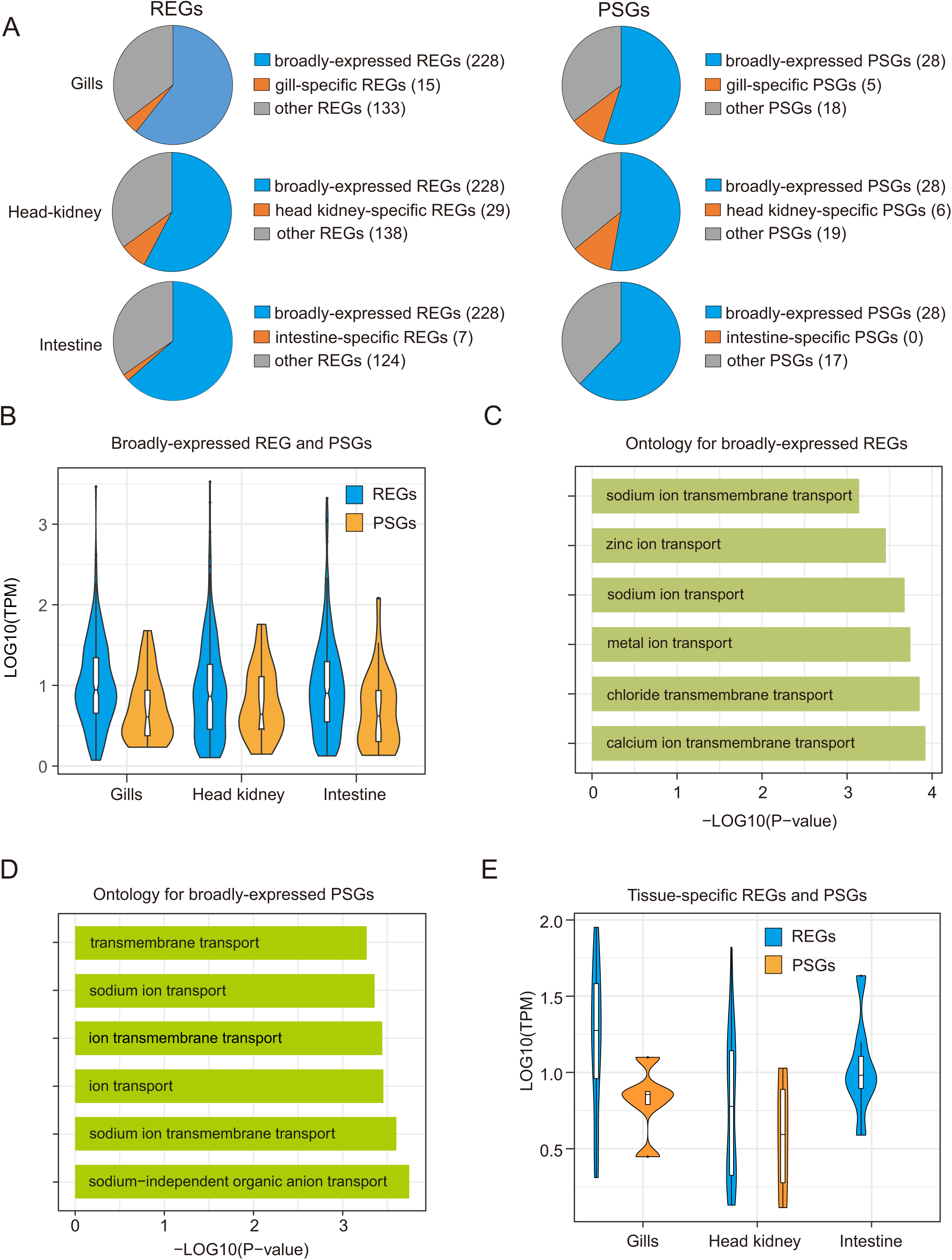
Expression patterns of REGs and PSGs in the three tissues of *G. p. kelukehuensis*. (A) Distribution of broadly-expressed, tissue-specific and other REGs and PSGs in each tissue, including gills, head kidney and intestine. (B) Violin plot depicting the distribution of Log_10_(TPM) values of broadly-expressed genes labeled by REGs and PSGs in three tissues. (C) Bar plot depicting the top six gene ontology for broadly-expressed REGs. Ontology for broadly-expressed REGs involved in ion transport functions. (D) Bar plot depicting the top six gene ontology for broadly-expressed PSGs. (E) Violin plot depicting the expression ranges of tissue-specific REGs and PSGs.

Interestingly, tissue-specific REGs and PSGs exhibited significantly different expression patterns. Although few tissue-specific REGs were identified in each tissues, we found that more tissue-specific REGs expressed in head kidney than other tissues (FIG. 3A, supplementary table S11). In addition, very few tissues-specific PSGs were identified, and no intestine-specific PSGs was determined (FIG. 3A, supplementary table S12). Among the tissue-specific REGs, we found gill-specific REGs had higher expression levels than head kidney-specific and intestine-specific REGs (FIG. 3E). Moreover, we failed to identify ion transport-related tissue-specific REGs or PSGs in *G. p. kelukehuensis*.

### Transport-associated genes co-expressed involved in alkaline tolerance

After a weighted gene co-expression network analysis (WGCNA) based pairwise correlations between gene expression trends across all samples, we ultimately obtained nine modules, and genes within modules were highly interconnected (supplementary table S13). A set of genes in nine modules exhibited distinct signature of expression patterns in all three tissues, that is said, these genes were interacted (supplementary table S14, FIG. S1). We found both dark green and royal blue modules showed highest and significant correlation to three tissues in *G. p. kelukehuensis*. After filtration and selection of modules including REGs or PSGs, we found royal blue module included 12 REGs and one PSGs. Specifically, this set of genes were enriched in ion transport and energy metabolism (FIG. 4).

**FIG. 4.**
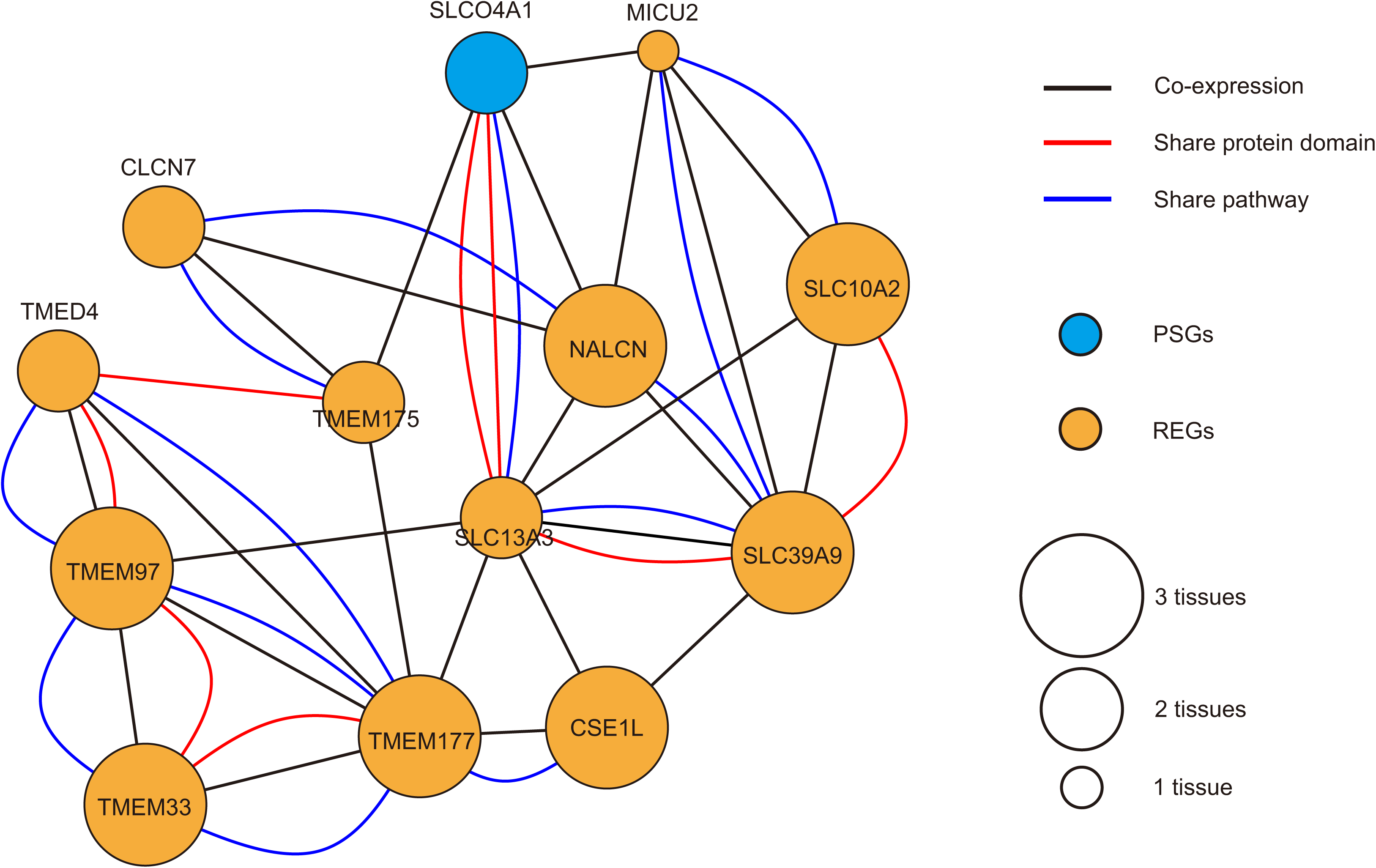
Network of co-expressed REGs and PSGs involved in alkaline tolerance. The colored line indicated the type of connections between genes. The colored solid circles indicated REGs and PSGs. The hollow circle indicated the number of tissues.

## Discussion

The increasing of water salinization is a growing threat to freshwater fish species (Herbert et al. 2015; Hintz & Relyea 2017). Endemic to the Tibetan Plateau, the Schizothoracinae (Teleostei: Cyprinidae) is the largest and most diverse taxon of the Tibetan Plateau ichthyofauna (Wu 1992; Cao et al. 1980). As the Tibetan highland lakes are long suffering gradually increased water salinization, unexceptionally, Schizothoracine fishes are forced to adapt to ongoing environmental changes (Tong, Fei, et al. 2017; Zheng 1997; Jiang et al. 2007). Comparative genomics has begun to elucidate the genomic basis of harsh environment adaptation in Tibetan fish species, but insight into the alkaline adaptation in Schizothoracine fishes has lagged behind. Building on previous comparative genomic or transcriptomic studies specifically focused on hypoxia and cold adaptation of Tibetan fishes (Yang et al. 2014; Chi et al. 2017; Kang et al. 2017; Wang et al. 2015; Zhang et al. 2017; Tong, Tian, et al. 2017; Ma et al. 2015), we performed one of the first comparative genomics studies to identify the signature of ongoing alkaline adaptation in Schizothoracine fishes on the Tibetan Plateau. These putative genomic signatures included expansions of lineage-specific genes associated with ion transport and transmembrane functions, genome-wide elevated rate of molecular evolution in Schizothoracine fishes relative to other model teleost fish species. In addition, we found specific changes in the rate of molecular evolution between *G. p. kelukehuensis* and other teleost fishes for ion transport-related genes. Furthermore, we identified a set of genes associated with ion transport and energy metabolism underwent positive selection. Using tissue-transcriptomics, we found that most REGs and PSGs in *G. p. kelukehuensis* were broadly expressed across three tissues and significantly enriched for ion transport function. Finally, we identified a set of ion transport-related genes with evidences for both selection and co-expressed which contributed to alkaline tolerance.

Polyploidization is a very common scenario found in teleost fishes (Leggatt & Iwama 2003; Van de Peer 2004). Noteworthy, most Schizothoracine fish species are polyploidy, such as tetraploid, and even sixteen-ploid (Wu 1992), the high complexity and large size of ployploid fish genomes has constrained the development of genomic resources for Schizothoracinae. As the recent advances in sequencing technologies, the genomes of several polyploidy fish species had been sequenced (Xu et al. 2014; Lien et al. 2016; Van de Peer 2004), including a recently available draft genome of a Schizothoracine fish, *Oxygymnocypris stewartii* (Liu et al. 2019). Given that, transcriptomics is a rapid and effective approach to obtain massive protein-coding genes for comparative genomics analysis among Schizothoracine fishes (Tong, Tian, et al. 2017; Tong, Fei, et al. 2017; Yang et al. 2014). Unlike seven diploid teleost fish with whole genomes, *G. p. kelukehuensis* is tetraploid. The completeness of transcriptome assembly (BUSCO) and genome content comparison results both suggested that gene models of *G. p. kelukehuensis* were similar to those representative well-annotated teleosts. In addition, we generated more than 6,000 pairwise orthologous gene groups between *G. p. kelukehuensis* and other teleost fish genomes, and identified over 2,000 putative single-copy orthogs among Schizothoracine fishes. The genome-wide phylogeny based on thousands of orthologs pinpointed the phylogenetic position of *G. p. kelukehuensis* within Schizothoracinae clade, which supported the previous phylogeny based on mitochondrial markers (He & Chen 2007). This finding emphasizes the application of transcriptome for further large-scale comparative genomics in Schizothoracine fishes.

We found an elevated rate of molecular evolution (dN/dS) in Schizothoracinae clade including *G. p. kelukehuensis* lineage on both concatenation and coalescent genome-scales, indicated that *G. p. kelukehuensis* may be under accelerated evolution. This finding is consistent with previous comparative genomics studies compared dN/dS ratios for very small scale of Schizothoracinae species (Tong, Tian, et al. 2017; Yang et al. 2014; Tong, Fei, et al. 2017; Chi et al. 2017). Similarly, previous genome-wide studies also suggested highland animals underwent accelerated evolution relative to lowland ones (Qiu et al. 2012; Qu et al. 2013). Studies very often interpret high dN/dS as a putative sign of positive selection (Bakewell et al. 2007). Species shard similar ecological niches can be shaped by convergent evolution to form physiological or morphological similarities (Stern 2013). Like other Tibetan terrestrial animals, our finding suggested that elevation of genome-wide molecular evolutionary rates is one of strategies that Schizothoracine fish including *G. p. kelukehuensis* adapted to the extreme environment of the Tibetan Plateau, such as the increasing of water salinization.

Lineage-specific gene families are sets of paralogs expanded and often exist in one specific lineage or species (Lespinet et al. 2002). We identified a large number of lineage-specific genes associated with ion transport processes in *G. p. kelukehuensis*, such as water transport, response to pH and monovalent inorganic cation transport, indicating this set of genes showed the putative signature of gene family expansion. This result was consistent with findings in Amur ide that dwelt in an alkaline environment in Lake Dali Nur (Xu et al. 2017) and another Schizothoracine fish living in saline lake Qinghai (Tong, Fei, et al. 2017), indicating that the alkaline environment of Lake Keluku, Lake Dali Nur and Lake Qinghai spurred evolution and expansion of genes in ion transport function. In this study, we identified a bunch of both REGs and PSGs repertoires involved in ion transport process in *G. p. kelukehuensis*, including the members from solute carrier (SLC) family and transmembrane protein (TMEM) family. This finding is similar to the recent studies of species dwelt in alkaline environment (Xu et al. 2017; Tong, Fei, et al. 2017). SLC is a super family that encoded transmembrane transporters for inorganic ions, amino acids, neurotransmitters, sugars, purines and fatty acids, and other solute substrates (Dorwart et al. 2008). Recent studies suggested that the adaptive evolution of SLC genes could contribute to saline and alkaline tolerance in fishes (Xu et al. 2017; Dorwart et al. 2008; Kavembe et al. 2015). Such as SLC4 subfamily, it encodes bicarbonate-transporter and regulated of Cl^−^-HCO_3_^−^ exchange and plays significant roles in maintenance of intracellular pH equilibrium (Pushkin & Kurtz 2006; Alper 2006). The SLC13 genes are the family of sodium sulphate/carboxylate cotransporters (Markovich & Murer 2004). In addition, the H^+^/Cl^-^ exchange transporter (CLC) genes mediate transmembrane Cl^−^ transport (Basilio et al. 2014). Altogether, we suggested that ion transport related genes underwent rapidly evolving or positive selection are another signature of alkaline adaptation in *G. p. kelukehuensis*.

Besides the genes involved in ion transport functions, we also identified a set of energy metabolism related genes under positive selection in *G. p. kelukehuensis*, especially related to mitochondria synthesis. A number of previous genome-wide studies on Tibetan terrestrial animals and several fishes suggested that an increased evolutionary rate and positive selection on genes involved in energy metabolism, which contributed to highland adaptation (Qiu et al. 2012; Qu et al. 2013; Yang et al. 2014; Wang et al. 2015). Although our study mainly focused on alkaline adaptation in Schizothoracine fish, similar to other Tibetan native animals, *G. p. kelukehuensis* had to adapt the long-term low temperature aquatic environment on the Tibetan Plateau. In addition, massive transport processes need continuous supplement. This finding suggested that *G. p. kelukehuensis* had developed strong capacity to meet high-energy demands in adaptation to chronic cold and alkaline stress.

Fish gills, kidney and intestine are three key tissues contributed to alkaline tolerance (Wilkie & Wood 1996; Heath 2018). To gain insight into the expression pattern and interaction of genes under selection, which contributed to ongoing alkaline adaptation, we characterized the expression profiles of three tissues types (gills, head kidney and intestine) and constructed the regulation network in *G. p. kelukehuensis*. Most of REGs and PSGs were broadly expressed across all three tissues, indicating that genes experiencing accelerated evolution in *G. p. kelukehuensis* may have general functions, and involved in multiple biological processes. Notably, broadly expressed REGs and PSGs mainly involved into sodium ion transport, chloride transport and metal ion transport functions. In addition, past studies emphasized that transmembrane protein (TMEM) and solute carriers (SLC) were broadly expressed in multiple tissues and played roles in biological processes (Dorwart et al. 2008; Markovich & Murer 2004; Tusnády et al. 2004). In this study, we first constructed the co-expression network of transport-related genes under selection in Schizothoracine fish, indicating that genes with similar functions and interact by cooperation in alkaline adaptation. Therefore, a more thorough understanding of interactome contributed alkaline adaptation in Schizothoracine fish can be achieved by increasing sequencing and function assay.

## Date Archiving

The Illumina sequencing reads have been deposited at NCBI Sequence Read Archive under the NCBI BioProject accession PRJNA577103.

## Acknowledgements

This work was supported by grants from the National Natural Science Foundation of China (31870365).

